# A Kmer-based paired-end read (KPR) *de novo* assembler and genotyper to genotype major histocompatibility complex class I (MHC-I) alleles for the dog

**DOI:** 10.1101/2020.07.15.205559

**Authors:** Yuan Feng, William H. Hildebrand, Stephen M. Tompkins, Shaying Zhao

## Abstract

The major histocompatibility complex class I (MHC-I) genes are highly polymorphic among individuals. MHC-I genotyping is required for determining the antigen-binding specificity of each MHC-I molecule in an individual. Numerous tools have been developed for human MHC-I genotyping using deep sequencing data such as RNA-seq; however they do not work for the dog, due to very limited information for canine alleles. To address this issue, we developed a Kmer-based paired-end read (KPR) *de novo* assembler and genotyper, which first assemble paired-end RNA-seq reads mapped to the MHC-I regions into contigs *de novo* and then genotype each contig. Our KPR tools are validated by Sanger sequencing, simulation and published genotype data. Applying our KPR tools on the published RNA-seq data of 158 tumor and 64 normal samples from 158 dogs, we have achieved a genotyping success rate of 86%, which includes 133 tumor and 57 normal samples from 142 dogs. We have identified 39 known alleles and 83 new alleles of high confidence in these dogs, yielding a more comprehensive MHC-I allele diversity landscape for the dog.

## Introduction

A human individual expresses 3-6 alleles encoded by three classical MHC-I genes HLA-A, HLA-B and HLA-C (human MHC-I will thus be referred to as HLA-I hereafter). Canine MHC-I genes include DLA-88, DLA-12 and DLA-64 (canine MHC-I will be referred to as DLA-I hereafter). However, only DLA-88, being the most diversified and with the highest expression level, is considered a classical MHC-I gene^1-4^. The status of DLA-12 and DLA-64 is currently unclear^1^, due to the limited information for these loci.

The three MHC-I genes share >90% sequence identity overall. Moreover, their exons 2 and 3, which encode the antigen-binding pocket, are among the most polymorphic regions in the genome. MHC-I genotyping (accurate determination of the about 550bp sequence of exons 2 and 3 of each allele) of an individual is required for determining the binding of each MHC-I molecule to an antigen. Thus, MHC-I genotyping is important for research on cancer (e.g., tumor specific neoantigen discovery), infectious disease, organ transplantation and other health issues.

MHC-I genotyping is traditionally achieved by PCR-based Sanger sequencing^1-4^. While considered the gold standard, this method is time-consuming, labor-intensive and low-throughput^5^. As numerous individuals have undergone deep sequencing (e.g., RNA-seq for gene expression analysis), many algorithms are developed for MHC-I genotyping using these massive data^5-12^. The challenge is that these sequence reads, generated by Illumina or similar next– generation sequencing (NGS) technologies, are often short (usually <150bp). This, coupled with the high sequence homology among the three MHC-I genes and their highly polymorphic nature among individuals, complicate the genotyping.

In humans, the challenge is addressed by taking advantage of the vast number of known HLA-I alleles^5-14^, about 19,180 in the current IMGT/HLA database^15^. With the assumption that many HLA-I alleles existing in the human population are already defined^15^, the current HLA-I genotyping tools^5-13^ focus on accurately mapping the NGS reads to the known allele reference, and then genotyping the individual by identifying known alleles with the most unambiguously placed NGS reads. Even for a few tools that perform assembly, the goal is either to simply extend the short reads somewhat to achieve more accurate mapping to known alleles^10^, or the assembly is reference-based but not *de novo*^*14*^. Hence, these assembly-performing tools are still heavily relying on known alleles.

In dogs, known DLA-I alleles are very limited, with only 95 total (76 alleles for DLA-88, 16 alleles for DLA-12, and 3 alleles for DLA-64)^1-4^. Hence, current MHC-I genotyping tools^5-14^., whose accuracy heavily depends on the comprehensiveness of known alleles, do not work for the dog. To address this deficiency, we are developing new software tools, as described below.

## Methods

### Kmer-based paired-end read (KPR) *de novo* assembler and genotyper

Our KPR assembler assembles DLA-I alleles *de novo*, as illustrated Figure 1A and below.

- Step 1: DLA-I read-pairs are identified by first mapping paired-end RNA-seq reads of a dog, after cleaning with Trimmomatic^16^ (version 0.36), to the DLA-I reference, which consists 103 known DLA-I alleles (having official allele names^4^) and 50 alleles derived from GenBank^1,4,17,18^ (no official allele name assigned yet). Then, coordinately mapped pairs with at least one read placed to exon 2 or 3 are identified, and further selected with each base quality at ≥30 to reduce sequencing error rate to ≤0.1%.
- Step 2: K*me*r dictionary is built from the forward (or reverse) read of each pair selected above at an exhaustive 1bp sliding window. Low frequency Kmers are be identified, and read-pairs harboring any of them are removed to further reduce sequencing errors.
- Step 3: Read pair extension starts with a randomly chosen pair with forward read *F* and reverse read *R*. It is extended by another pair with forward read *f*_i_ and reverse read *r*_*i*_, if *F* and *R* share identical sequence of >K in length with *f*_*i*_ and *r*_*i*_, respectively, at their 3’ or 5’-end (Figure 2A). This yields *F*_*contig*,_ and *R*_*contig*_, which are then looped through the same extension process until no further extension is possible.
- Step 4: Final contig is assembled from the last *F*_*contig*,_ and *R*_*contig*_ from step 3, if they share identical sequence of >K at their 5’ or 3’-end.
- Step 5: Steps 3 and 4 are repeating *N* (e.g., >30,000) times such that all extension paths have been exhausted.

Our genotyper genotypes assembled contigs, as illustrated in Figure 1B and described below.

- Contig sorting and variant linkage building: Assembled contigs are classified as normal, or as DLA-88*50X group (50X)^1^ which has a 3-base insertion at a specific location in exon 3. Multiple sequence alignment of each group yields a consensus sequence with variable sites and variant linkage groups identified.
- Variant linkage validation and frequency estimation with paired-end reads: To reduce false results from misassembly, variant linkage groups are validated with paired-end reads (as the two sequences of a pair are from the same allele), after mapping the DLA-I read-pairs (see step 1 in the assembly section) to the contigs. The frequency of each linkage group is estimated with read-pair coverage at its unique linkage sites.
- Contig extension (if not completely covering exons 2 and 3): This is because MHC-I genotyping requires the entire exon 2 and 3 sequence. Incomplete contigs are first extended by other incomplete contigs via overlapping variant linkage or paired-end reads (Fig 2B). If still incomplete, these contigs are further extended with paired-end reads based on variants or frequency (Fig 2B).
- DLA-I genotyping: Each complete contig (with entire exons 2 and 3) is translated into amino acid sequence, and compared to the known DLA-I reference for gene and allele identification. Allele group will be assigned by examining the sequences from three hypervariable regions (HVRs), following the scheme by the International Society for Animal Genetics DLA Nomenclature Committee^4,17^, including 3-digit typing for allele group (e.g., DLA-88*034), and 5-digit typing for allele (e.g., DLA-88*034:01).

**Figure 1.**
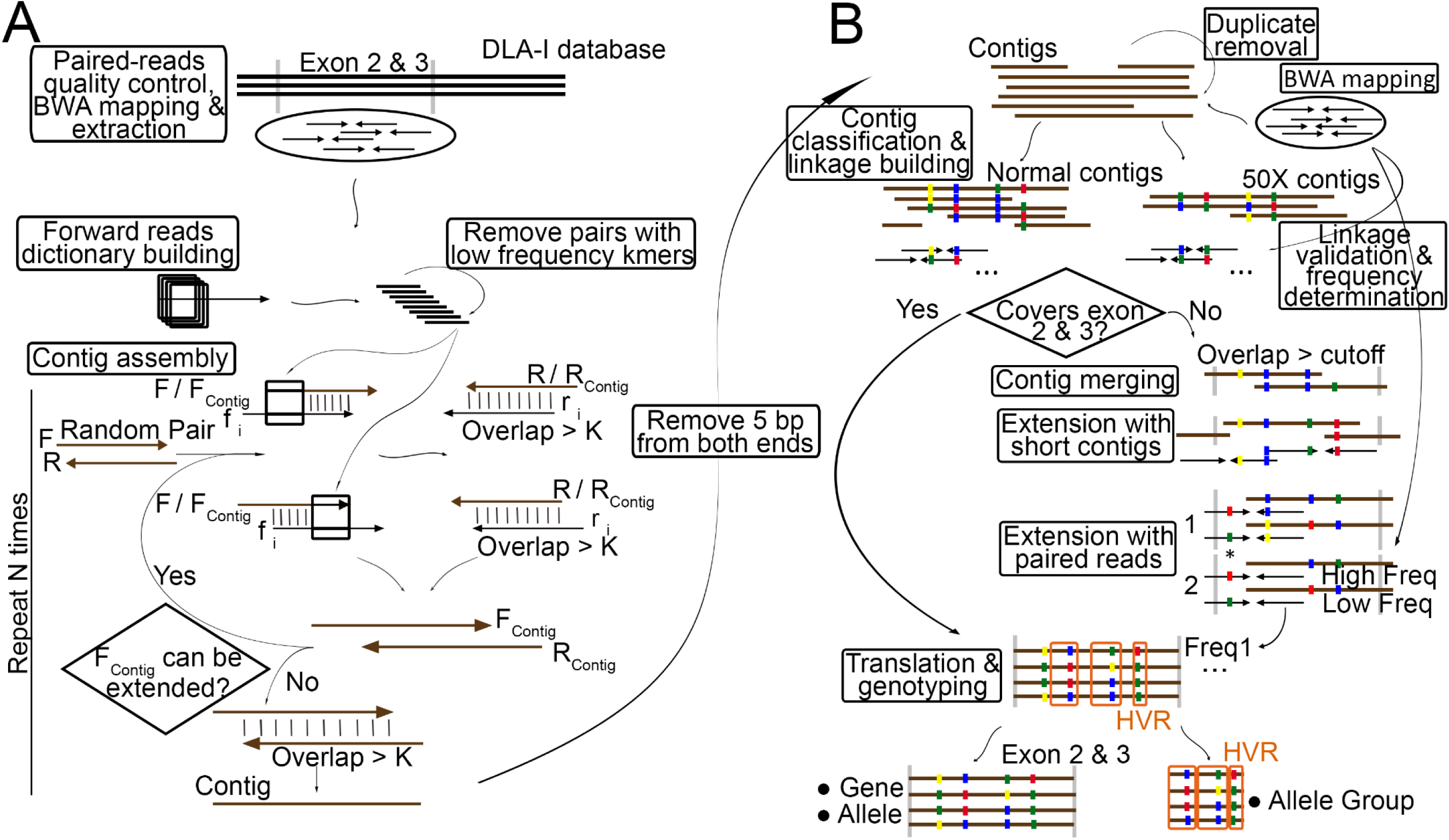
Our tools rely on paired-end sequencing to genotyping MHC-I alleles. A. Our KPR assembler assembles the highly polymorphic region, the entirety of exons 2 and 3, of DLA-I alleles of an individual *de novo*, using paired-end RNA-seq reads. Contigs are represented by bars, while paired-end reads are represented by paired-arrows facing each other. B. Our genotyper genotypes each assembled contig after variant linkage building, validation and if needed, extension. Sequence variants are indicated by colored dots.

**Figure 2.**
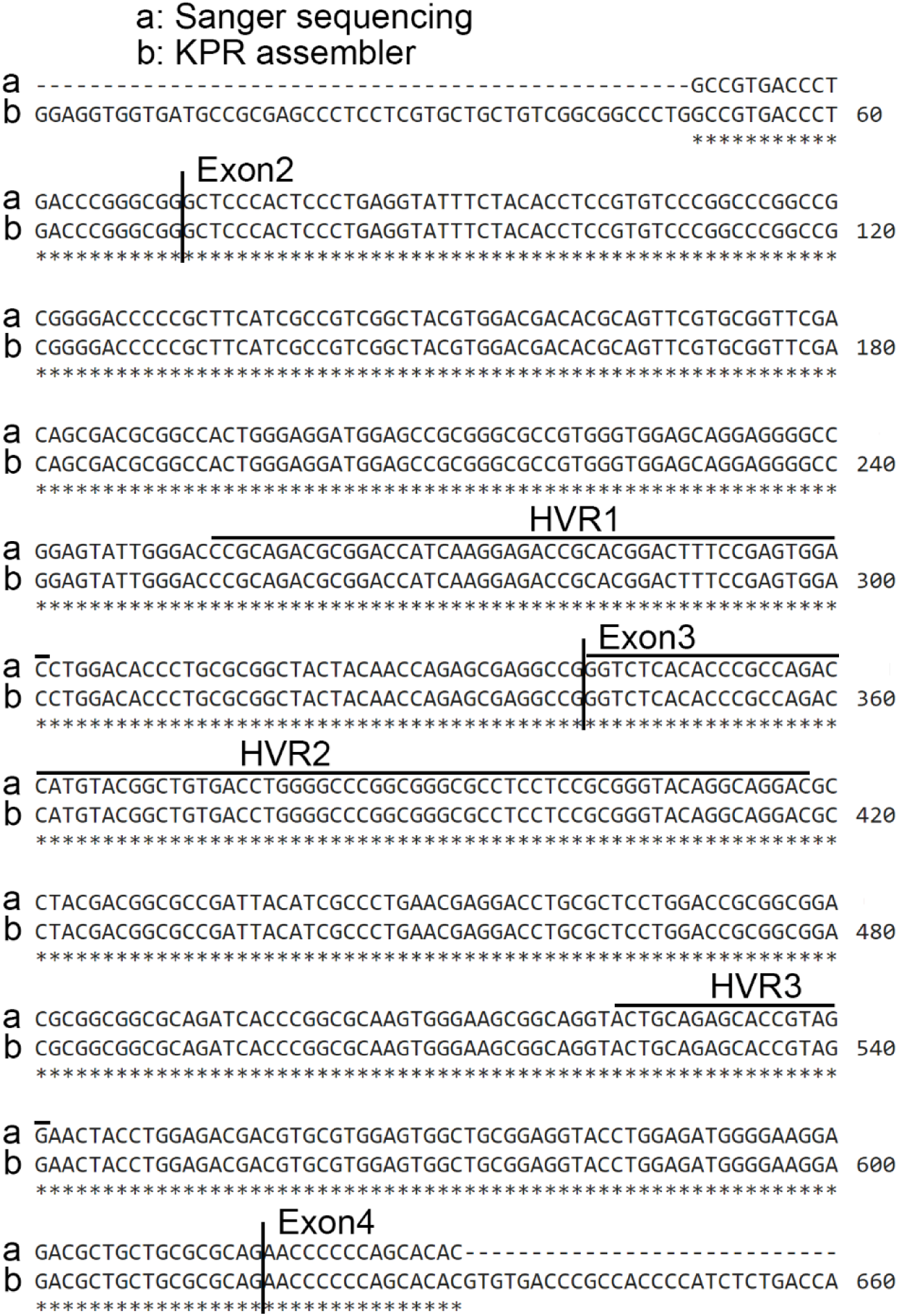
Contigs assembled by our tools are validated by Sanger sequencing. Sanger sequence is identical to the assembled sequence obtained by applying our KPR tools to the RNA-seq data of a dog. HVRs: hypervariable regions reported previouly^17^

### Sanger sequencing

Sanger sequencing was performed as described^3^. Exons 2 and 3 of DLA-88 and DLA-12 were amplified from cDNA samples of 5 English Cocker Spaniel dogs, as well as tumor and/or normal samples of 5 other dogs with mammary cancer^19^. The primers used are “GGGAGAGAGTCCAGGGTAGG” (forward) and “GGTTTCACTTCTGCGTCTCC” (reverse) for DLA-88, and “CGACCCTAAAGGTCTGGGCTA” (forward) and “ACCACTGGCGGTTATCTCAG” (reverse) for DLA-12 (Table S2A), designed by Primer 3^20^.

### Sample simulation

Six paired tumor and normal RNA-seq samples from 3 dogs from a published study^21^ were used for the simulation. They are: 1) SRR7779554 and SRR7779476 from dog CMT-162; 2) SRR7779670 and SRR7779671 from dog CMT-785; 3) SRR7779469 and SRR7779468 from dog CMT-149 (Table S3B). Via random sampling of the DLA-I database, 8 alleles combinations were made to simulate all likely scenarios, including homozygosity versus heterozygosity, and DLA-88*normal versus DLA-88*50X. The 8 combinations are listed in Table S3A. As a result, a total of 6 x 8 = 48 samples were simulated.

Simulated DLA-I read pairs of each sample was generated from selected known alleles, considering 3 parameters derived from the real RNA-seq data of the sample. First, sequencing errors were estimated by aligning the RNA-seq reads to the canFam3 reference genome with BWA^22^ (version 0.7.17) and then counting the number of mismatches. Sequencing error rates of the six samples range from 0.331% to 0.483%, with which sequencing errors were introduced to the simulated RNA-seq reads. Second, based on the mapping position to the known alleles in the SAM files, each simulated read-pair was made to replace its corresponding real RNA-seq read pair, having the same read length and insert size. Third, the expression levels of the 3 DLA-I genes were estimated via a unique region in Exon 3 (“ACCATGTAC” for DLA-88, “TGGATGTTT” for DLA-12, and “TGGACTTCG” for DLA-64). Based on this, the number of simulated read-pairs for each DLA-I allele was determined.

### Positive predictive value (PPV) and root mean square error (RMSE)

PPV was calculated by 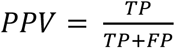, using allele frequencies to represent each positive call.

RMSE was calculated by 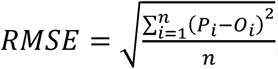, where *P*_*i*_ represents the observed frequency of allele *i*, while *O*_*i*_ represents the corresponding ground truth frequency.

### Genotyped 158 dogs with published RNA-seq data^21^

RNA-seq data of 158 tumors and 64 normal samples (222 total) of 158 dogs (Table S4A) published^21^ were downloaded from the SRA database (PRJNA489087), and quality controlled as follows. First, MultiQC^23^ (version 1.5) was used to examine GC content and duplicate level (Figure S4B-C, S4F). Base quality distribution before and after Trimmomatic trimming was also examined (Figure S4A). Second, tumor-normal sample pairing accuracy was evaluated using germline variants (Figure S4D). Third, breed validation was performed with using breed-specific germline variants.

After QC, each sample was then genotyped using our KPR *de novo* assembler and genotyper, as described before and outlined in Figure 1.

### RNA-seq genotyping validation with whole-exome sequencing data

Whole-exome sequencing (WES) data from the same 158 dogs genotyped was downloaded from the SRA database (PRJNA489159). Germline variants were used to validate the pairing of WES and RNA-seq samples via dog IDs (Figure S4E).

Validation with WES read pairs: WES read pairs of a sample were first cleaned by Trimmomatic with default setting, and then mapped to the reference, which consists of all RNA-seq-based contigs, with BWA. All concordantly mapped reads, including both uniquely and repeatedly mapped, were extracted. For each read, its best matched RNA-seq contig was identified. Finally, an RNA-seq contig is validated if it has the most or the 2^nd^ most WES reads from the same dog mapped (Figure S4G).

Validation with WES contigs: The assembly was conducted separately for exon 2 and exon 3, following the step outlined in Figure 1. WES contigs were then compared with RNA-seq contigs with BLAST. An RNA-seq contig is validated if it is matched with an identity >95% and an E-score <1e-5 by a WES contig that covers >85% of exon 2 or 3 and is from the same dog.

### Allele clustering and bootstrapping

DLA-I alleles were clustered with their sequence identities determined by Clustal-Omega multiple sequence alignment of either complete exon 2 and 3 sequences, or pseudo-sequences generated with HVRs. Heatmaps and dendrograms were created by “gplot” in R. Euclidean distance and complete agglomeration method were used in both sequence clustering and bootstrapping.

### HVR refinement

Shannon diversity index, Simpson diversity index and DIVAA^24^ were used to identify potential HVRs in DLA-I molecules. Only “validated” alleles (see Results) were used for the analysis. Three cutoffs were used for HVR identification: 1) minimum amino acid diversity; 2) minimum length of the region; and 3) minimum number of positions with required diversity within the region (Figure S6A-C).

Human MHC-I alleles were downloaded from the IPD and IMGT/HLA database^25^, including 5012 HLA-A, 6326 HLA-B and 4923 HLA-C alleles.

## Results

### Contigs assembled by our tools are validated by Sanger sequencing

We applied our KPR assembler (Figure 1) to our canine RNA-seq data, 27 to 87 million paired-end reads of 76×76bp or 50×50bp each of dogs with various cancers^19,26-28^. We then performed Sanger sequencing on 13 samples to validate the assembled contigs. The results (Tables S2B and S2C), with an example shown in Figure 2, indicates that our tools are effective.

### Our tools are optimized and validated by simulation

For our simulation, we chose six paired normal and tumor samples from three dogs, with one well-genotyped by our tools and the other having issues, from a published study^21^. For the well-genotyped dog, our tool identified a dominant allele, DLA88*012:01, in both tumor and normal samples, with 80% and 73% allele frequency respectively (Table S3B). For those with issues, our tool found completely different alleles between the tumor and normal samples of one dog, and more than the expected numbers (3 in tumor and 8 in normal samples) of DLA-88 alleles in each sample of another dog (Table S3B).

For our first simulation which aims to optimize the running parameters, we randomly chose 1 or 2 alleles per DLA-I gene from the reference allele database. For each allele, we simulated its read-pairs for each of the six samples described above, based on the expression level of the corresponding DLA-I gene (Figure S3A), read length, insert-size and sequencing errors determined with the actual RNA-seq data of the sample. Then, we replaced the real DLA-I read pairs with simulated read-pairs. We made a total of 8 allele combinations per sample, by varying DLA-88 allele type (normal or 50X) and homozygosity or heterozygosity of each DLA-I gene (Table S3A). This yields a total of 6 x 8=48 simulated samples.

We genotyped the 48 simulated samples by varying running parameters K (the Kmer length) and N (the number of assembly runs) (Figure 1), as well as sequencing depth D. We used two values, positive prediction value (PPV), which measures the genotyping accuracy without considering the genotyped allele frequency, and root mean squared error (RMSE), which measures the difference between the ground truth and the genotyped allele frequency, to evaluate the performance of our tools.

We first set *N* = 30,000 and kept the original D, and genotyped each sample by changing K from 20mer (unique in a mammalian genome) to 80mer (as the maximum read length is 101bp^21^). The median PPV increases to 1 (100% accuracy) when K is between 40bp and 70bp (Figure 3A), while the median RMSE is the lowest when K is 50bp (Figure 3A). The results indicate that the optimized K is 50bp for running our tools for these samples. Second, by setting *K* = 50*bp* and keeping the original D, we varied N from 100 to 70,000. Both PPV and RSME indicate that the genotyping results are stabilized when N is ≥30,000 (Figure 3B). Third, we set *K* = 50*bp* and *N* = 30,000, and varied D from 1 million (M) to 100M pairs. The analysis was separated for the 3 DLA-I genes, because their expression levels differ. The results indicate that the minimum total amount of RNA-seq read-pairs required for reliable assembly is 15M for DLA-88, 30M for DLA-12, and 100M for DLA-64 (Figure 3C).

**Figure 3.**
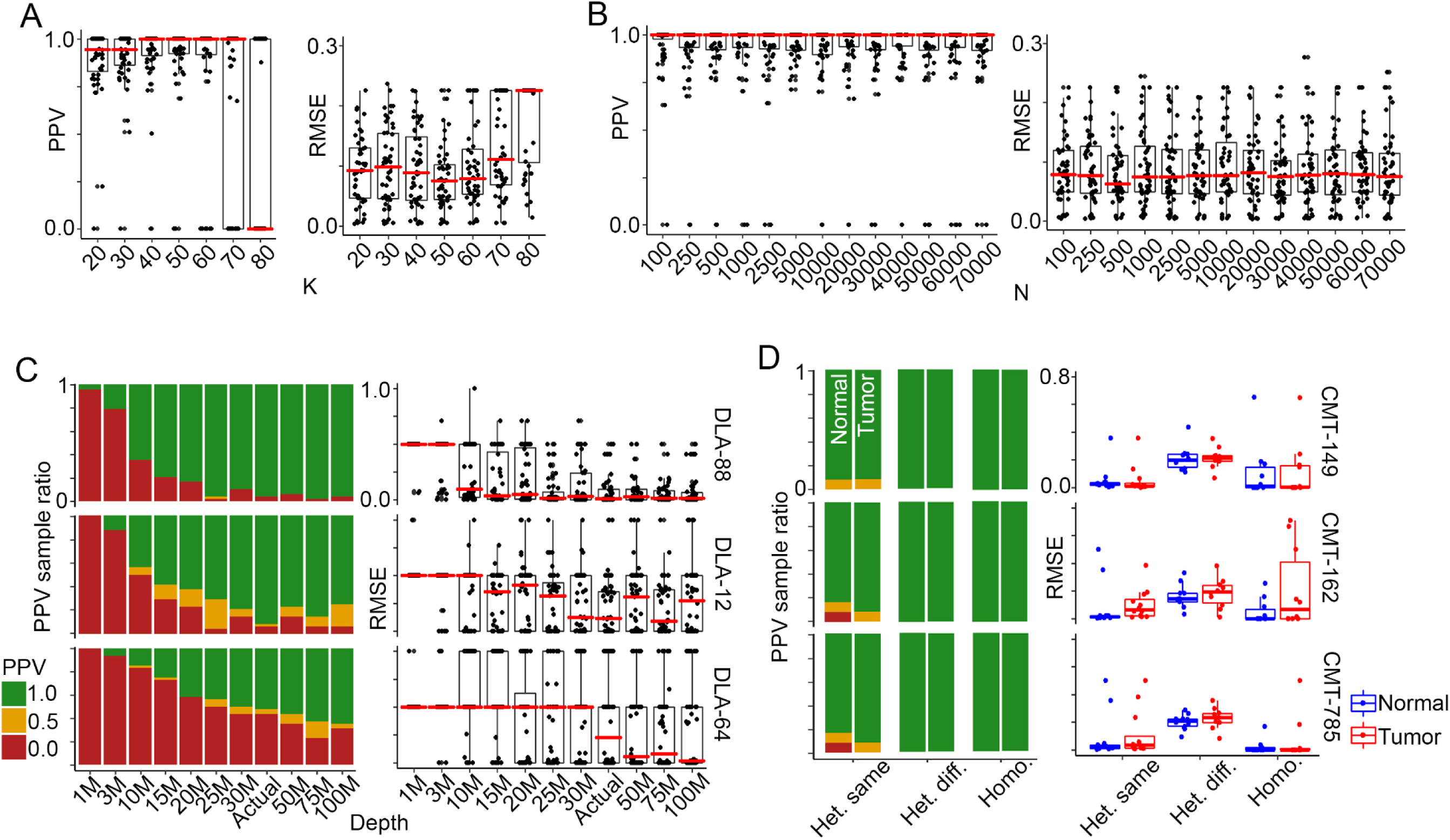
Our tools are optimized and validated by simulation. A-B. Parameter optimization. Varying K (Kmer length) and N (the number of assembly runs) are evaluated via positive predictive value (PPV) and root mean square error (RMSE), indicating that the optimal values are K = 50 and N >30,000. C. Determination of the minimum sequencing depth, D, for DLA-I genotyping. The 3 DLA-I genes were independently evaluated, and the minimum depths for effective DLA-88, DLA-12 and DLA-64 genotyping are 15M, 30M, and 100M, respectively. D. The assembly of DLA-88 with different allele combinations. Homozygous alleles are better performed, compared to heterozygous alleles of the same group (either normal or 50X), or heterozygous alleles of different groups. No significant differences are observed between normal and tumor samples.

Interestingly, we did not observe any significant differences in the simulation between the well-genotyped dog and dogs with genotyping issues (Table S3B). To determine if this is allele-related, we expanded our simulation by randomly selecting alleles for each of the 8 allele combinations (Table S3A), and studied individual DLA-I genes in each sample (Figure 3D). As expected, DLA-88 is well genotyped, while DLA-64 is not. However, the analysis again finds no significant differences among the three dogs (Figure 3D). The reasons may be as follow.

First, for the dog where the tumor and normal samples have different genotyping results (Table S3B), the samples may express different DLA-I genes. For the dog with >2 DLA-88 alleles genotyped per sample, sequencing errors, mis-assemblies due to more distant alleles (Figure S3C), or sample contamination with another dog may be the issues.

Besides parameter optimization, the simulation results also indicate our KPR assembler is effective. This is because at the optimized conditions (K = 50bp; N ≥ 30,000; D ≥ 30M), our KPR tools consistently achieve a genotyping accuracy of *PPV* = 1 and *median RMSE* < 0.1 for DLA-88 (Figure 3).

### We genotyped 158 dogs with an 86% success rate and discovered 83 new DLA-I alleles of high confidence

These 158 dogs are from a mammary cancer study^21^. RNA-seq data, 25 to 77 million read pairs of 101×101bp per sample, are published for their tumors and in 64 dogs, also matching normal samples^21^ (Table S4A), yielding 222 samples in total.

We performed quality control (QC) of the data. First, we examined sequencing amount, base quality, read duplication and GC content (Figure S4A-C, S4F). Then, we used germline mutations discovered in each sample to validate the tumor-normal pairing accuracy (Figure S4D), as well as breed data accuracy (Figure S4E). We found that two Yorkshire Terrier dogs are actually Shih Tzu (Figure S4E), and made the correction. Other than this, all samples passed our QC measures.

We applied our KPR tools on the 222 samples of 158 dogs. We genotyped 190 samples (133 for tumors and 57 for normal) from 142 dogs, achieving a success rate of 86% (32 samples failed genotyping, as no complete contigs of exon 2 and 3 sequence were assembled). We identified 39 known alleles and 105 new allele candidates in the genotyped dogs (Figure 4A). The results indicate our KPR tools are effective, as described below.

**Figure 4.**
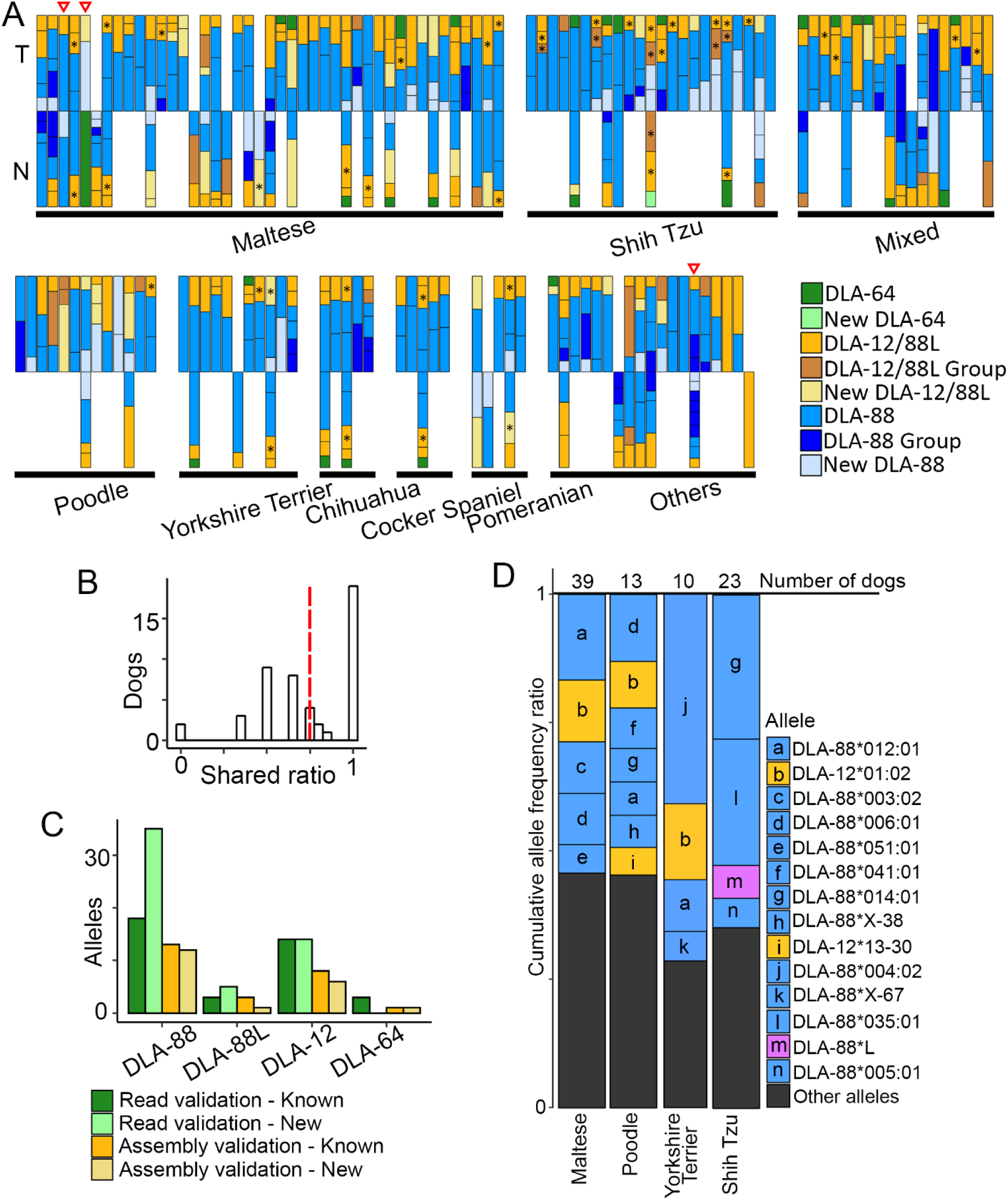
We successfully genotyped 142 dogs and discovered 83 new DLA-I alleles of high confidence. A. The DLA-I genotyping results of 142 dogs obtained by applying our KPR tools to the paired-end RNA-seq reads of their tumor (T), and normal (N) samples. A dog is represented by single or paired vertical bars. The lines inside each bar separate individual alleles, with the height indicating the estimated allele frequency. Asterisks denotes DLA-88L alleles. Red triangles indicate dogs used for the simulation analysis described in Figure 3. B. The distribution of agreement in genotyped alleles between tumor and normal samples of the same dogs. Vertical red dashed line indicates the median of agreement at 75%. C. Validation RNA-seq genotyping by WES read pairs or assemblies, with 85% known alleles and 57% new alleles validated. D. Breed-dominant alleles in 4 pure breeds with ≥10 dogs per breed. Only tumor samples are used for this analysis. The height of the blocks indicates the cumulative allele frequencies of the specific allele across all samples. Top 4 alleles or top alleles with combined frequency reaching >50% are shown.

First, consistent with literature reports^1-4^, DLA-88 is the most expressed. It is identified in 186 out of 190 samples in total, with combined allele frequency ranging from 8% to 100% and having a mean of 70% (Figure 4A; Table S4A). DLA-64 is the least expressed, found in 29 samples with a median frequency at 10% (Figure 4A; Table S4A). DLA-12 is expressed 147 samples with a median frequency at 17% (Figure 4A; Table S4A).

Second, using an allele frequency cutoff of 8%, we identified 1-2 alleles per gene in 101 out of our 142 dogs (Figure 4A; Table S4A). For the small number of dogs with >2 alleles per gene, likely reasons include contigs assembled because of sequencing errors, and the DLA-12 locus that may encode DLA-88 like alleles (to be discussed in later sections).

Third, consistent alleles between tumor and normal samples are identified for many dogs. Indeed, among 47 dogs where both tumor and normal samples are genotyped, close to half share identical alleles and only 2 dogs have no shared alleles (Figure 4B). One likely reason for disagreement may be that tumor and normal samples express different alleles.

Forth, we validated our RNA-seq genotyping results with WES data that are also published for these dogs^21^, using two approaches. First, we mapped WES read pairs to the RNA-seq assembled alleles (Table S4B). For each WES sample, we identified the top two RNA-seq alleles that have the most WES reads concordantly placed. Then, we determined if the WES sample and the two RNA-seq alleles are from the same dog. If yes, the allele is validated. We achieved a validation rate of 83% for known alleles and of 50% for new alleles (Figure 4C; Table S4D). Second, we also conducted assembly with WES data for exon 2 and exon 3 separately, and compared the resulting contigs with RNA-seq alleles from the same dog. As a result, 45 alleles were validated (Figure 4C). Combining both approaches, a total 93 alleles are validated with WES data, including 60 new alleles.

Lastly, consistent with published work^1,3^, our results indicate dominant alleles within a pure breed (Figure 4D). Specifically, we identified dominantly expressed alleles in 4 canine breeds with ≥10 dogs (Figure 4D; Figure S4H). Certain dominant alleles have a frequency of >25%, such as DLA-88*004:02 in Yorkshire Terrier, as well as DLA-88*014:01 and DLA-88*035:01 in Shih Tzu (Figure 4D). These Shih Tzu alleles match published studies^1^, and differ more from those of three other breeds (Figure 4D), consistent with their places of origin.

In summary, we have used our tools to successfully genotype these dogs. Importantly, we have discovered 83 new DLA-I alleles that are: 1) validated by WES data as described above (Figure 4C); 2) recurrent in ≥2 dogs; and/or 3) with ≥50% allele frequency in an RNA-seq sample.

Hence, they are less likely to be artifacts arisen from sequencing errors or mis-assembly, and are high confidence alleles.

### Three DLA-I genes differ in expression and diversity

With 39 known and 83 high confidence DLA-I alleles, we investigated their expression among the 142 dogs (Figure 4A). Consistent with previous studies that classify DLA-88 a classical MHC-I gene^1,4^, multiple DLA-88 alleles are highly expressed in numerous dogs (Figure 5A; Figure S5A). With only one allele expressing in a few dogs (Figure 5A; Figure S5A), our study supports that DLA-64 is not a classical MHC-I gene.

**Figure 5.**
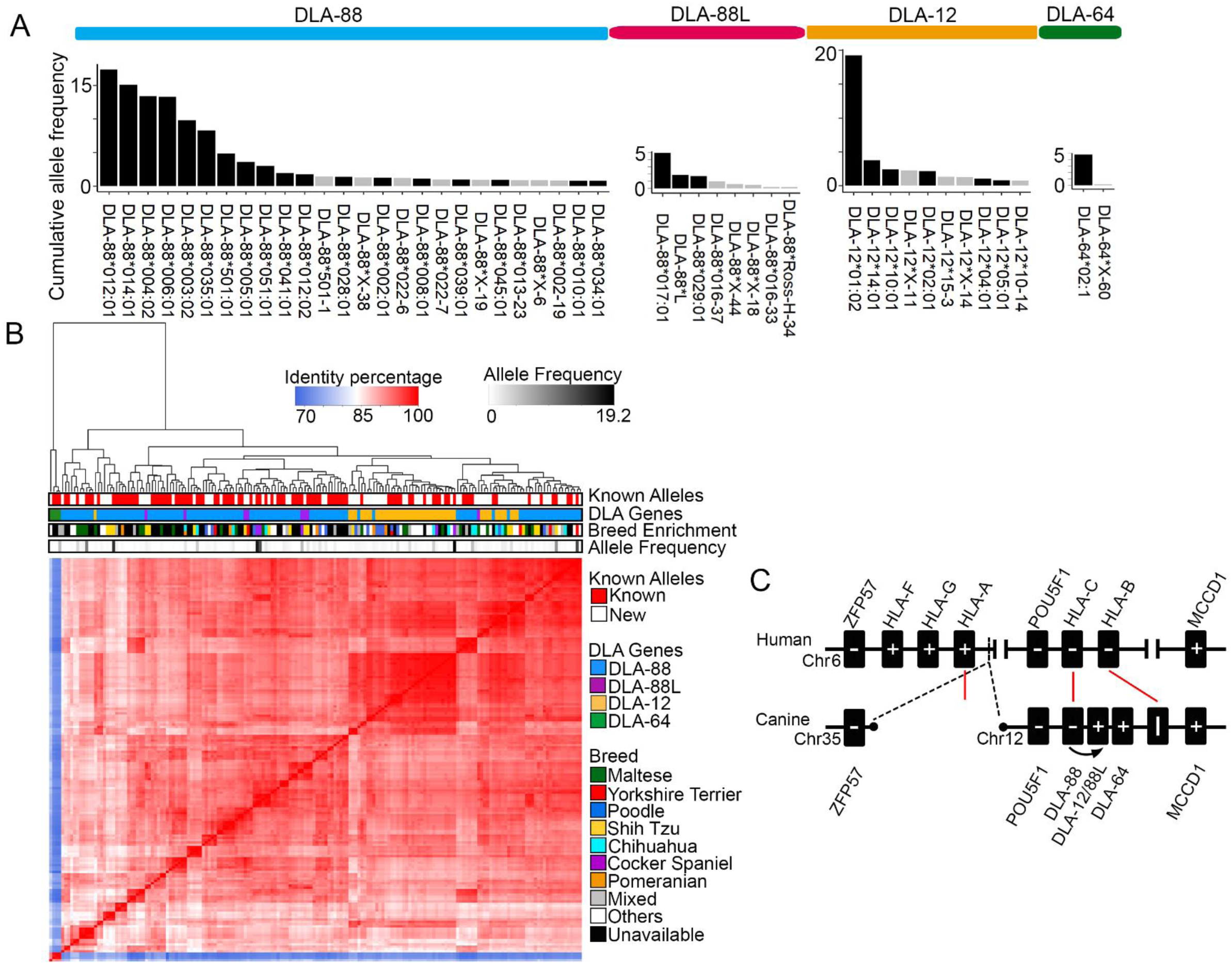
Three DLA-I genes differ in expression and diversity. A. Distribution of cumulative allele frequency in 142 dogs examined. For DLA-88 and DLA-12, only alleles with cumulative allele frequency of >0.7 are shown, while all DLA-64 and DLA-88 alleles are shown. B. The clustering of known (95 alleles) and new alleles of high confidence (83). Three groups are clearly observed: 1) a DLA-64 group; 2) a distant DLA-88 group; and 3) a DLA-88, DLA-88L and DLA-12 group. The 3^rd^ group is further clustered into DLA-88, DLA-88/DLA-88L and DLA-88/DLA-12 subgroups. Breed enrichment score represents the fraction of dogs within a breed that carry the allele. C. Proposed evolution of DLA-I genes. Red lines link the corresponding HLA-I and DLA-I genes. Dash lines indicate the breakage of the HLA-I locus in the canine genome. “-” and “+” specify the gene orientations.

DLA-12 is more complicated, as a previous study indicates that the locus actually expresses two genes DLA-12 and DLA-88L^1^. We found that DLA-12 and DLA-88L express in these dogs with allele diversity and expression level between DLA-88 and DLA-64 (Figure 5A; Figure S5A).

We performed clustering analysis with amino acid sequence alignments among all 95 known alleles and 83 high confidence alleles, 178 in total (Figure 5B). As expected, DLA-64 alleles are outliers (Figure 5B). Interestingly, DLA-88 alleles are divided into several clusters, with one clearly distinct and perhaps more ancient (Figure 5B) because of the enrichment in Shih Tzu and Maltase^29^. The remaining cluster can be further divided into 3 subgroups described as DLA-88, DLA-88/DLA-88L, and DLA-88/DLA-12 (Figure 5B). While Yorkshire Terrier dogs are found in both DLA-88/DLA-88L and DLA-88/DLA-12 subgroups, Poodle dogs mostly locate in the DLA-88/DLA-12 subgroup (Figure 5B). These results indicate that DLA-88L and DLA-12 are distinct allele groups (Figure 5B). DLA-88L appears to be found in only certain breeds, including Shih Tzu, consistent with the previous study^1^. We also observed possible “pioneers” in these major clusters, which have significantly higher allele frequencies compared to other members in the cluster, indicating a potential “founder” effect (Figure 5B).

We investigated the dog-human synteny of the MHC-I loci, and found that DLA-88 is clearly the homologue of HLA-C gene (Figure 5C). The canine homologue of HLA-A is possibly lost when the ancestral region of the HLA-locus broke in the canine genome, forming the end of canine chromosome 35 (Figure 5C). At the canine orthologous site of HLA-B, we only find a pseudogene (Figure 5C); thus it is conceivable that the canine homolog of HLA-B is also destroyed. Because of the highly homology of DLA-12/88L to DLA-88 (Figure 5B), we hypothesize that the DLA-12/88L locus is duplicated from the DLA-88 site (Figure 5C). We do not know the origin of DLA-64.

### We further refined HVRs in DLA-I alleles

Three HVRs have been previously reported for DLA-88, and are used to assign the allele groups^4^. With our 52 newly identified DLA-88 (47 alleles) and DLA-88L (5) alleles of high confidence, combined with 76 known alleles in the database, we reexamined HVRs via rigorous statistical analyses. Briefly, we determined the amino acid diversity at each position by computing DIVAA^24^, Shannon, or Simpson diversity index from the alignment of 128 alleles (Table S6). To discover HVRs, we first determined the cutoff of amino acid diversity at each individual site (Figure S6A), and identified the sites that meet the cutoff, which we call diversified sites. Then, by examining the distance distribution between two neighboring diversified sites, we determined a cutoff to segment DLA-I amino acid sequences, as shown in Figure S6B. Those segments with the diversified site content above a minimum cutoff (Figure S6C) are HVRs (Figure 6).

**Figure 6.**
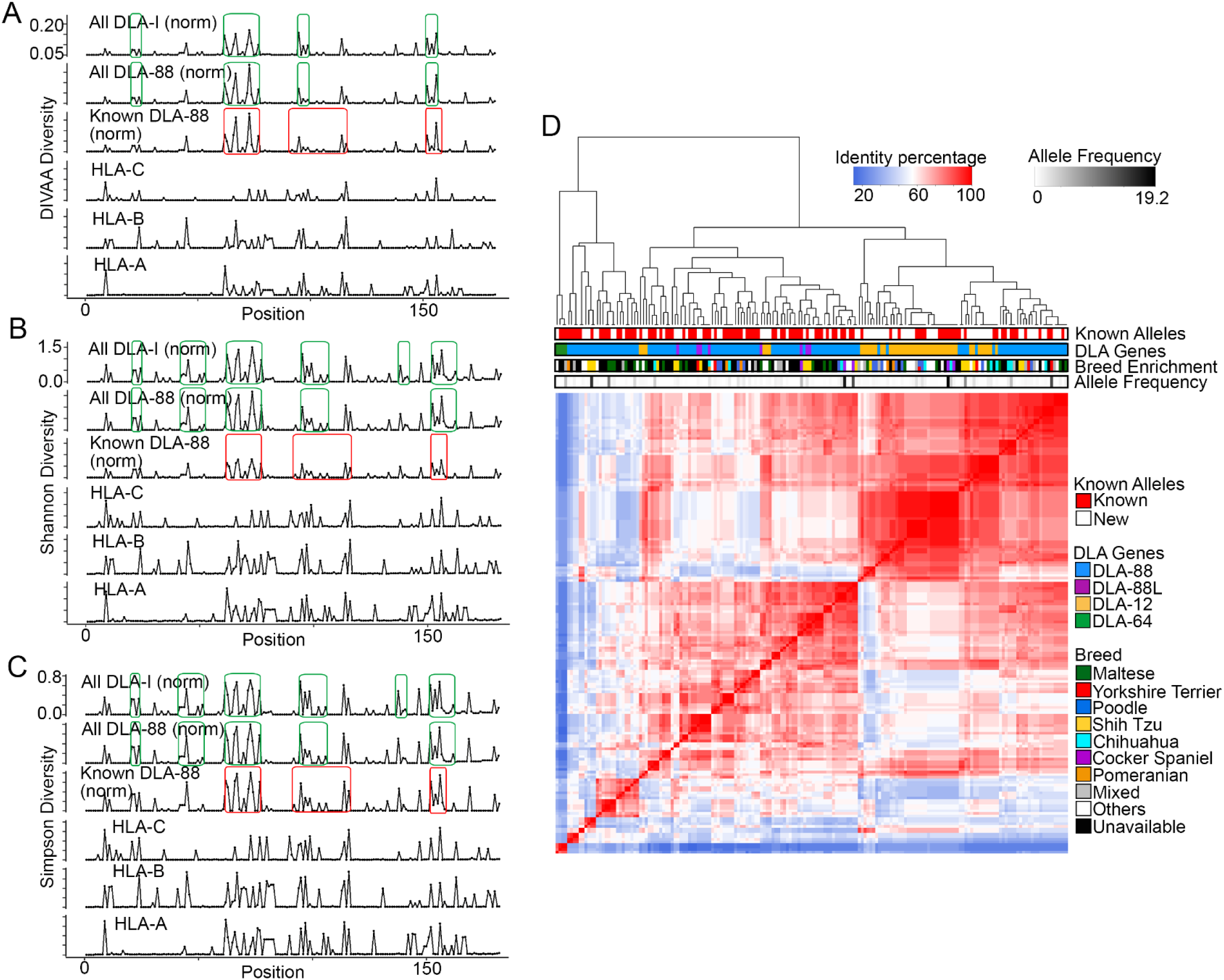
We further refined HVRs in DLA-88 alleles. A-C. HVRs are identified with DIVAA^24^, Shannon or Simpson diversity index among 178 DLA-I, 128 DLA-88, 76 known DLA-88 alleles, as well as among 5,012 HLA-A, 6,326 HLA-B and 4,923 HLA-C alleles respectively. For simplicity, DLA-88*50X alleles (with 183 amino acids) are excluded. Green boxes represent HVRs identified by us, while red boxes represent HVRs reported prveiouly^4,17^. D. Heatmap indicates the clustering of 178 DLA-I alleles with HVRs identified using DIVAA diversity index shown in panel A.

HVRs identified by the DIVAA^24^, Shannon, or Simpson diversity index largely agree (Figure 6A-C). Importantly, most of our newly identified HVRs overlap with those reported previously (Figure 6A-C). However, differences are also noted. First, we uncover a new HVR at the N-terminal site and significantly narrow down the 3^rd^ HVR (Figure 6). Lastly, the corresponding human regions of canine HVRs are also hyper-diversified among the three HLA-I genes (Figure 6).

We repeated the HVR discovery by including DLA-12 (30 new and 16 known) and DLA-64 (1 new and 3 known) alleles for 178 alleles in total. This however does not change the HVRs identified with the DIVAA index, but uncovers an extra 4 amino acid-HVR with the Shannon and Simpson’s diversity index.

To determine the contribution of HVRs to the four clusters identified in Figure 5B, we performed the same clustering analysis using HVR sequences only (Figure 6D). The same clusters are identified, indicating the identified HVRs truthfully reflecting the diversity of the DLA-I alleles.

## Discussion

### KPR *de novo* assembler and genotyper

The dog serves as an important translational model in various human diseases, including cancer, infectious disease, obesity, neurological and other disorders. The canine model has the great potential to effectively bridge a current gap between preclinical models and human clinical trials^30-35^, accelerating drug discovery. This is because, unlike rodent models, canine diseases arise spontaneously in animals with intact immune systems, thereby recapturing the essence of human diseases. As companion animals, dogs share the same environments as humans and are exposed to many of the same carcinogens. For example, environmental toxins, advancing age and obesity are known risk factors for cancer development in dogs^30^, the same as in humans.

Thus, creating resources that are critically missing at present, such as tools for canine MHC-I genotyping, is urgently needed.

In recent years, NGS-based MHC-I genotyping has become a focus and numerous software tools have been published^5-14^ for human HLA-I genotyping. Unfortunately, these tools, which are built upon the ∼19,180 human alleles, do not work for the dog. This is because only 95 canine alleles are known currently. We are trying to address this issue by developing new software tools, the KPR *de novo* assembler and genotyper, reported here.

Because our core algorithm relies on paired-end sequencing (note that paired-reads are both derived from the same allele), the accuracy of our tools is not as heavily influenced by known alleles as are existing methods^5-14^. Indeed, Sanger sequencing validation, simulation analysis, and consistence with published results^1^ all indicate that our tools are effective.

However, our tools needs improvement. First, for a dataset with 158 dogs, the success rate of our tools is ∼86%. The reason for failed genotyping is the assembled contigs are incomplete, not containing the entire exon 2 and 3 sequence. To address this issue, we plan to implement dynamic programming, such as the Floyd-Warshall algorithm, in the assembly process. We have successfully implemented dynamic programming in our copy number software SEG^36^. Second, our current tools are designed for RNA-seq, but not for WGS and WES data, which contain intronic sequences. Importantly, unlike RNA-seq, WGS and WES lack sequence reads that directly link exons 2 and 3. Thus, new functionality is needed.

### DLA-I allele diversity landscape

We have used our KPR tools to successfully genotype 142 dogs with public RNA-seq data^21^, which leads to the discovery of 83 new alleles of high confidence and dominant alleles in 4 pure breeds. Notably, our clustering analyses with both known and new alleles consistently reveal four groups among DLA-88 alleles: 1) a distant and perhaps more ancient DLA-88 group; 2) DLA-88 group; 3) DLA-88L containing group; and 4) DLA-12 containing group. Our findings indicate that DLA-88L and DLA-12 alleles are closer to certain DLA-88 alleles, compared to more distant DLA-88 alleles themselves. A published study reports that the DLA-12 locus encodes DLA-12 in 80% of dogs and a DLA-88L (DLA-88 like) gene in 20% of dogs examined^1^. Another study reports >2 alleles of DLA-88 in individual dogs^3^. These results support that the DLA-12 locus is duplicated from the DLA-88 locus.

Our analysis indicates that DLA-88 is homologue of HLA-C. However, we have not found the canine homologue of either HLA-A or HLA-B. We do not know if they are lost during evolution, or simply missing because the genome assembly is incomplete for these sites, which are near the chromosomal end. We have also searched other assembled canine genomes, including the genomes of Great Dane and Basenji, but have not found the homologues either. Future research is needed to clarify this issue.

Finally, adding the newly found DLA-I alleles, we have refined their HVRs, which are currently used for DLA-88 allele group classification^4^. However, we note that HVRs also harbor sites that are not diversified, and that certain highly diversified sites locate outside HVRs. Hence, we propose that using “hyper-variable sites” (HVSs) (Figure S6G) may better capture the diversity of DLA-I alleles, leading to more accurate classification of DLA-I alleles and prediction of the DLA-I/antigen binding specificity.

## Acknowledgments

We thank Joshua Watson and Maxwell Colonna for editing the manuscript; and the Georgia Advance Computing Resource Center (GACRC) for supporting this work. This work is funded by NCI R01 CA182093.

